# Genomic analysis of *Staphylococcus aureus* isolates from bacteremia reveals genetic features associated with the COVID-19 pandemic

**DOI:** 10.1101/2023.12.22.572975

**Authors:** Miquel Sánchez-Osuna, Marc Pedrosa, Paula Bierge, Inmaculada Gómez-Sánchez, Marina Alguacil-Guillén, Mateu Espasa, Ivan Erill, Oriol Gasch, Oscar Q. Pich

**Author notes:** Address correspondence to Miquel Sánchez-Osuna or Oscar Q. Pich.

## Abstract

Genomic analyses of bacterial isolates are necessary to monitor the prevalence of antibiotic resistance genes and virulence determinants. Herein, we provide a comprehensive genomic description of a collection of 339 *Staphylococcus aureus* strains isolated from patients with bacteremia between 2014 and 2022. Nosocomial acquisition accounted for 56.6% of episodes, with vascular catheters being the predominant source of infection (31.8%). Cases of fatality (27.4%), persistent bacteremia (19.5%) and diagnosis of septic emboli (24.2%) were documented. During the COVID-19 pandemic, we observed a 140% increase of the episodes of *S. aureus* bacteremia per year, with a concomitant increase of the cases from nosocomial origin. This prompted us to investigate the existence of genetic features associated with *S. aureus* isolates from the COVID-19 pandemic. While genes conferring resistance to β-lactams (*blaI-blaR-blaZ*), macrolides (*ermA, ermC, ermT, mphC, msrA*) and aminoglycosides (*ant*(4’)-Ia, *ant*(9)-Ia, *aph*(3’)-IIIa, *aph*(2’’)-Ih) were prevalent in our collection, detection of the *msrA* and *mphC* genes increased significantly in pandemic *S. aureus* isolates. Similarly, we observed a higher prevalence of isolates carrying the genes encoding the Clumping Factors A and B, involved in fibrinogen binding. Of note, macrolides were extensively used as accessory therapy for COVID-19 and fibrinogen levels were usually elevated upon SARS-CoV-2 infection. Therefore, our results reveal a remarkable adaptation of the *S. aureus* isolates to the COVID-19 pandemic context and demonstrates the potential of whole-genome sequencing to conduct molecular epidemiology studies.

## INTRODUCTION

Bacteremia is a life-threatening condition with high morbidity and mortality rates that can elicit a systemic host response known as sepsis (Viscoli 2016). These episodes often result in complicated bacteremia, usually defined by the presence of attributable mortality, the development of hematogenous embolisms (i.e. endocarditis, discitis, osteomyelitis of systemic abscesses) or the persistence of viable bacteria in blood after three or more days after proper antibiotic treatment (Gudiol et al. 2015). *Staphylococcus aureus* is among the top species causing bacteremia-associated mortality (Kern and Rieg 2020) and persistent bacteremia (Wiggers et al. 2016), which in turn is linked to a higher risk of metastatic spread (Khatib et al. 2006). There is increasing awareness that patients who survive sepsis often have long-term physical, psychological and cognitive disabilities with significant health care and social implications (Singer et al. 2016).

Several host factors such as certain comorbidities, the setting of the infection onset and the severity of sepsis, have been associated with the high mortality rates observed in *S. aureus* bacteremia (van Hal et al. 2012; Yilmaz et al. 2016). However, the patient’s age is the most consistent predictor of mortality, with older patients being twice as likely to die (van Hal et al. 2012). Also, certain strain characteristics such as methicillin resistance have also been shown to impact the clinical outcome (Lee et al. 2018). In contrast, the role of specific genetic traits of *S. aureus* in the evolution of bacteremia is much less understood. Recent studies suggest that *S. aureus* genetic factors predictive of infection evolution may be lineage specific, which would hinder their identification (Gasch et al. 2013; Recker et al. 2017).

*S. aureus* has also been described as a leading cause of secondary infection during past viral pandemics, significantly increasing patient mortality rates (Chertow and Memoli 2013). In this regard, recent reports have noted a higher incidence of bacteremia by *S. aureus* at the beginning of the COVID-19 pandemic (Falces-Romero et al. 2023). In the same line, our group has recently reported a rise of vascular catheter-related bacteremia in early 2020. Moreover, we also observed a significant increase in cases of *S. aureus* bacteremia (Gasch et al. 2022). However, while several studies have addressed the incidence, prevalence and clinical outcomes of SARS-CoV-2 and *S. aureus* co-infection (Adler et al. 2020; Adalbert et al. 2021), the genomic differences between such strains and those isolated in the pre-pandemic times have not been explored.

Here, we investigated the clinical aspects, epidemiology and genomic features of 339 *S. aureus* strains isolated from patients with bacteremia at the Parc Taulí University Hospital before and during the COVID-19 pandemic (2014-2022). This analysis allowed us to shed light into significant associations between *S. aureus* lineages, antibiotic resistance genes, virulence factors and the clinical outcome of bacteremia. Comparative genomics analysis pointed out that some virulence factors and antibiotic resistance genes were enriched in pandemic *S. aureus* isolates.

## MATERIAL AND METHODS

### Cohort and strain isolation

#### Patients and setting

All consecutive adult patients (≥18 years) diagnosed with *S. aureus* bloodstream infection from July 2014 to December 2022 at the Parc Taulí University Hospital (Sabadell, Spain) were retrospectively included in this study.

#### Clinical data

Clinical information was recorded including patient demographics, comorbidities and other clinical characteristics, source of bacteremia, place of acquisition, severity of sepsis, antibiotic therapy and infection outcomes. The modified Charlson score (Charlson et al. 1987) was used to assess patient morbidities. Setting of acquisition was classified according to modified Friedman criteria (Friedman et al. 2002) as: community, in-hospital acquired or healthcare-related. 30-day mortality was considered as any death within the first month after the bloodstream infection onset. Persistent bacteremia was defined as the presence of positive blood cultures after 72 hours of appropriate antibiotic therapy. Septic embolism was determined as the presence of one or more diagnosed secondary foci as a result of bacterial spread through blood. Strains isolated from March 14th 2020 (Spain’s lockdown start) onwards were considered as pandemic clones. These data are available in Data S1.

#### Strain isolation and manipulation

For each patient, only the first episode of *S. aureus* bacteremia was included in the analysis. Blood cultures were processed using the BACT/ALERT® automated system (bioMérieux) following standardized procedures. *S. aureus* strains were identified by using mass spectrometry technology (MALDI-TOF MS, Bruker). *S. aureus* single colonies were frozen and stored at -80°C. Antibiotic susceptibility testing was assessed by microdilution methodology according to standardized protocols using the MicroScan WalkAway system® (Beckman Coulter).

#### Ethics Statement

The Ethics Committee for Investigation with medicinal products (CEIm) of the Parc Taulí University Hospital approved the implementation of this study (2023/5088, approved on 6 October 2023). The requirement for informed written consent was waived given the retrospective nature of the study. Patient identification was encoded, complying with the requirements of the Spanish Organic Law on Data Protection 15/1999.

### Whole-genome sequencing and *de novo* assembly

#### DNA extraction

*S. aureus* isolates were grown on Columbia Agar with 5% Sheep Blood plates (bioMérieux) at 37°C. Total DNA was purified using the DNeasy Blood & Tissue Purification Kit (Qiagen) following the manufacturer’s instructions. Cell lysis was achieved by pre-incubating bacterial colonies in Phosphate-buffered saline (PBS) containing 100 μg/mL lysostaphin (Merck) at 37°C for 30 minutes. DNA quality was assessed using a NanoDrop device (Thermo Fisher Scientific) and a Qubit® 2.0 fluorometer (Thermo Fisher Scientific).

#### DNA sequencing

Libraries for sequencing were prepared with the Nextera XT DNA Sample Preparation Kit (Illumina). Whole-genome sequencing was performed using paired-end sequencing on an Illumina HiSeq 2500 and NovaSeq 600 sequencers available at the Genomics Unit of the Centre de Regulació Genòmica (CRG, Barcelona). Whole-genome sequencing data are available at National Center for Biotechnology Information (NCBI) under the accession number PRJNA1055690.

#### Quality control

The quality of the raw sequencing reads was checked with FastQC (Andrews 2010). Read pre-processing and filtering were performed with TrimGalore (Krueger 2012) by shaving the sequencing adapters, trimming the initial 20 bp poor-quality positions and using a Phred score >= 20 limit.

#### *De novo* assembly

The trimmed paired-end reads were assembled *de novo* using shovill (Seemann 2023) and annotated with prokka (Seemann 2014) against the COG (Galperin et al. 2021), HAMAP (Pedruzzi et al. 2015) and Pfam (Mistry et al. 2021) databases and using the *S. aureus* NCTC 8325 [NC_007795] proteome as the reference. CheckM2 was finally used for assessing the quality of assemblies (Chklovski et al. 2023).

### Isolate typing and *S. aureus* annotation

#### Isolate typing

Multilocus sequence typing (MLST) was carried out on the assembled scaffolds using the MLST software (Larsen et al. 2012). Sequence Types (ST) and Clonal Complexes (CC) were deduced from the *S. aureus* PubMLST database (Jolley et al. 2018) grouping the *arcC, aroE, glpF, gmk, pta, tpi* and *yqiL* gene types. spaTyper was used to assign *spa* types according to the Ridom Spa Server database guidelines (Sanchez-Herrero 2020), AgrVATE was used for *agr* typing (Raghuram et al. 2022) and SCCmec-types were obtained using SCCmecFinder (Kaya et al. 2018).

#### Antibiotic resistance genes and virulence factors

Putative antibiotic resistance genes (ARG) and virulence factors (VF) were predicted on the assembled scaffolds with ABRicate (Seemann 2017) using the in-built NCBI AMRFinderPlus database (Feldgarden et al. 2019) and Virulence Factor Database (VFDB) (Liu et al. 2019), respectively. Strains were considered as Panton-Valentine leukocidin (PVL) positive when both *lukF* and *lukS* genes were predicted on the same genome.

#### Plasmid prediction

Plasmid scaffolds were identified using MOB-recon and mobility was predicted based on the MOB-typer module (Robertson and Nash 2018). PlasmidFinder software from Center for Genomic Epidemiology (CGE) was used for *rep* typing (Carattoli et al. 2014) and roary for identifying plasmid core genes (Page et al. 2015).

#### Prophage prediction

Prophage prediction was performed as described recently for other *S. aureus* genomes (Sweet et al. 2023). Prophage regions were detected in assembled scaffolds using PhiSpy (Akhter et al. 2012) and CheckV was then used to remove host contamination and to delimit phage boundaries (Nayfach et al. 2021). Prophage clustering was carried out with usearch (Edgar 2010) so that sequences showing 90% similarity along their 90% length were counted as the same. Integrase detection and typing was assessed with BLASTP (Altschul et al. 1990) using previously reported integrase sequences (Goerke et al. 2009) as queries and limiting the e-value to 1e^−20^ and query coverage to >75%. Each putative integrase identified in the prophage regions was classified into an integrase group according to the query with the lowest e-value in the BLASTP analysis.

### Statistical methods

Mann-Whitney U (MWU) test was used for continuous variables and Fisher’s exact test (FT) for categorical variables. For clonal associations, MWU was first applied to determine biased distributions of lineages when analyzing a specific genetic feature. If the lineage distributions differed significantly when compared to global frequencies, FT was independently applied to determine the association of each feature with specific lineages. Clonal associations were only computed for strains with known lineage. Always a *p*-value threshold of 0.05 was considered as significant. Statistical analyses were performed using custom Python scripts.

## RESULTS

### Cohort description

We identified 339 consecutive adult patients diagnosed with *S. aureus* bloodstream infections between 2014 and 2022 (Data S1). Most of them were male (69.3%) and the mean age was 63.7 ± 19.9 years. The calculated Modified Charlson Score was greater or equal than 4 in 64.0%. The most frequent clinical conditions detected were diabetes (31.6%), the presence of a foreign device (25.4%), ischemic heart disease (17.7%), congestive heart failure (16.5%), solid neoplasm (15.3%), peripheral arteriopathy (12.7%) and chronic obstructive pulmonary disease (10.0%). Acquisition was nosocomial in 56.6%, while 27.4% were community-acquired and 16.0% healthcare related. Regarding the primary source of infection, vascular catheters (31.8%) and skin infections (18.0%) were predominant, followed by pneumonia (9.4%). As far as 30.7% episodes were of unknown origin (n=104). Antibiotic susceptibility tests indicated that 13.9% infections were caused by methicillin-resistant *S. aureus* (MRSA) strains (Data S1).

Regarding clinical outcomes, 93 cases were fatal (27.4%) and the median time to patient’s death was 20 (interquartile 25 (IQ25)=8, interquartile 75 (IQ75)=37) days (Data S1). Fatal bacteremia episodes showed to be significantly associated with age, modified Charlson, sepsis, congestive heart failure, immunosuppressive chemotherapy treatment, endocarditis and the presence of a cardiac device (Table S1). Of note, catheter-related bacteremia episodes were less lethal (*p* < 0.05) (Table S1). On the other hand, persistent bacteremia was observed in 19.5% of episodes, with the median duration being 43 (IQ25=24, IQ75=67.5) days (Data S1). Persistent infections were significantly associated with sepsis (Table S1). Pneumonia-originated bacteremia never resulted in persistent infections or septic emboli infections (*p* < 0.05) (Table S1). Septic emboli were diagnosed in 24.2% (Data S1) which were more prevalent in community-acquired infections (*p* < 0.05) (Table S1). Septic emboli infections were significantly associated with infective endocarditis episodes (Table S1).

### Isolate typing

The MLST software classified our *S. aureus* isolates in 24 different lineages (Figure 1A, Data S1). The major identified lineages were CC30 (19.5%), CC5 (17.1%), CC45 (12.1%), ST398 (8.8%), CC8 (8.8%), CC22 (8.0%) and CC15 (5.9%). 18 isolates could not be classified into a known sequence type. The distribution of lineages remained relatively stable over the years of study (Figure 1B).

**Figure 1.**
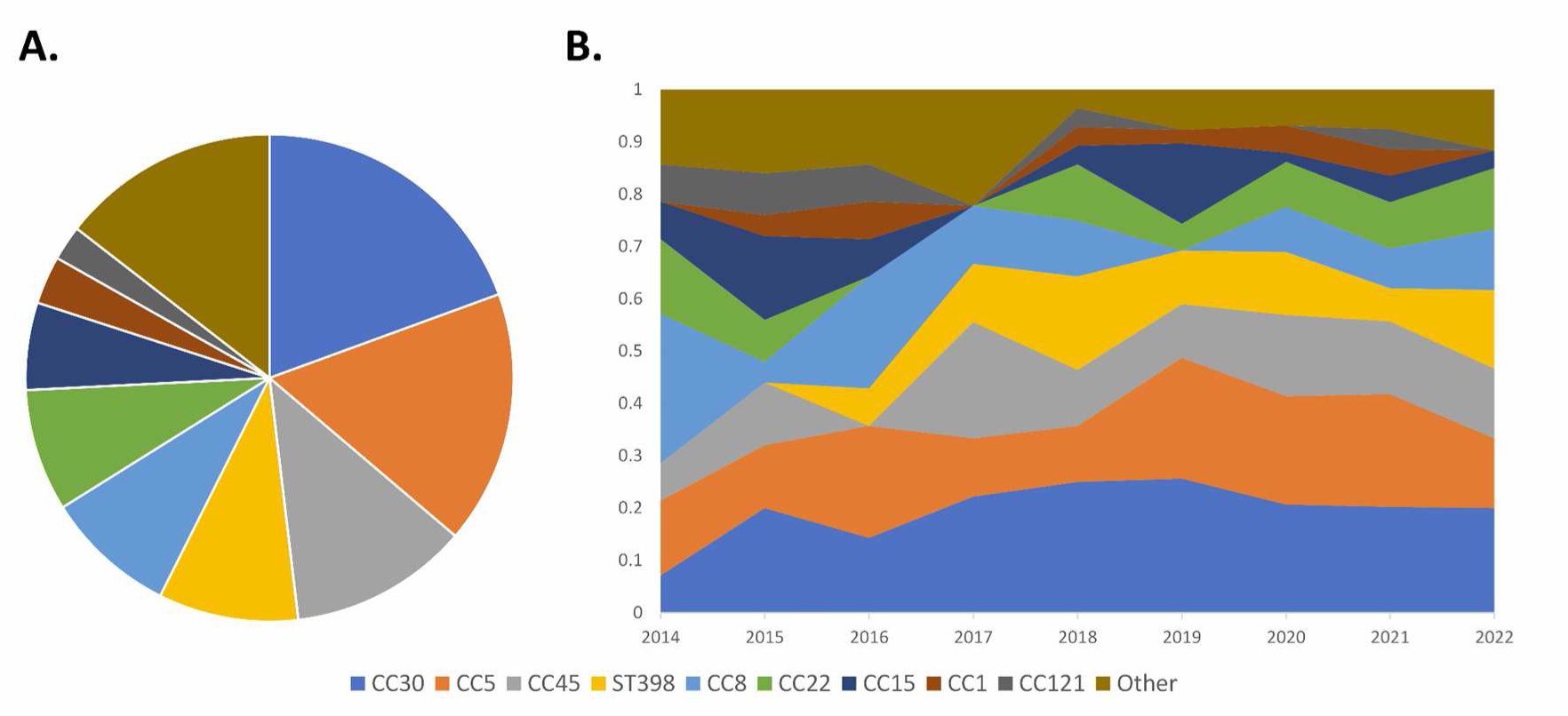
(A) Pie chart depicting global *S. aureus* lineage frequencies in our collection. (B) Graphical representation of *S. aureus* lineage frequencies over years. Lineages with less than 5 representatives are collected under the umbrella term “Other”.

We found significant associations between specific lineages and clinical data. CC5 and CC8 strains were more frequently recovered from patients with pneumonia-related bacteremia, while CC30 isolates were more prevalent among bacteremia from surgical site infections (*p* < 0.05) (Figure S1, Table S2). Although we did not find any significant association, CC1 and CC8 infections showed the highest mortality rates (45.4% and 40.0%, respectively). CC15, CC30, CC45, CC21 and ST398 lineages generated persistent infection with frequencies ranging from 20.0 to 25.0%, while CC1 (9.1%) and CC22 (7.4%) were the lineages generating fewer persistent infections. The strains most frequently associated with septic emboli were CC15 and CC45 (30.0% and 34.1%) while those that formed the least were CC1 and CC8 (9.1% and 10.0%) (Data S1).

MRSA strains belonged more frequently to CC5 (42.5%) and CC8 (38.3%) lineages (*p* < 0.05) (Figure S1, Table S2) but methicillin resistance was also found in CC22 (8.5%), CC45 (4.2%) strains and unknown lineages (6.5%) (Data S1). The vast majority (95.7%) of SCCmec cassettes were grouped into type IV, except for two SCCmec cassettes classified as type V (Data S1). SCCmec type IVc was associated with CC5 and CC8 while SCCmec type IV was linked to CC5 and CC22 clones (*p* < 0.05) (Figure S1, Table S2). In our collection, we did not find any association of MRSA strains with mortality (Matthews Correlation Coefficient (MCC) = 0.06).

Our analysis predicted that 13 isolates from our collection were *agr*-negative. Among the *agr*-positive strains, 48.5% were classified as type I, 26.1% type II, 22.1% type III and 3.3% type IV (Data S1). We observed a biased distribution of lineages regarding the *agr* types II and III (*p* < 0.05) (Figure S1). Specifically, we found that type II *agr* was associated with CC5 and CC15 isolates, while type III was almost exclusive of CC30 clones (*p* < 0.05) (Table S2). Finally, our typing analysis revealed 123 different *spa* types (Data S1), being t012 significantly associated with CC30 isolates, t002 and t067 with CC5, t008 with CC8, t084 with CC15 and t1451 with ST398 (Figure S1, Table S2).

### Antibiotic resistance genes

We identified homologs to 34 different ARG families conferring resistance to clinically relevant antibiotics (Figure 2, Data S1). Homologs of the *mecA* gene were found in 47 strains (13.9%) and they were associated primarily with CC5 and CC8 strains (*p* < 0.05) (Figure S1, Table S2).

**Figure 2.**
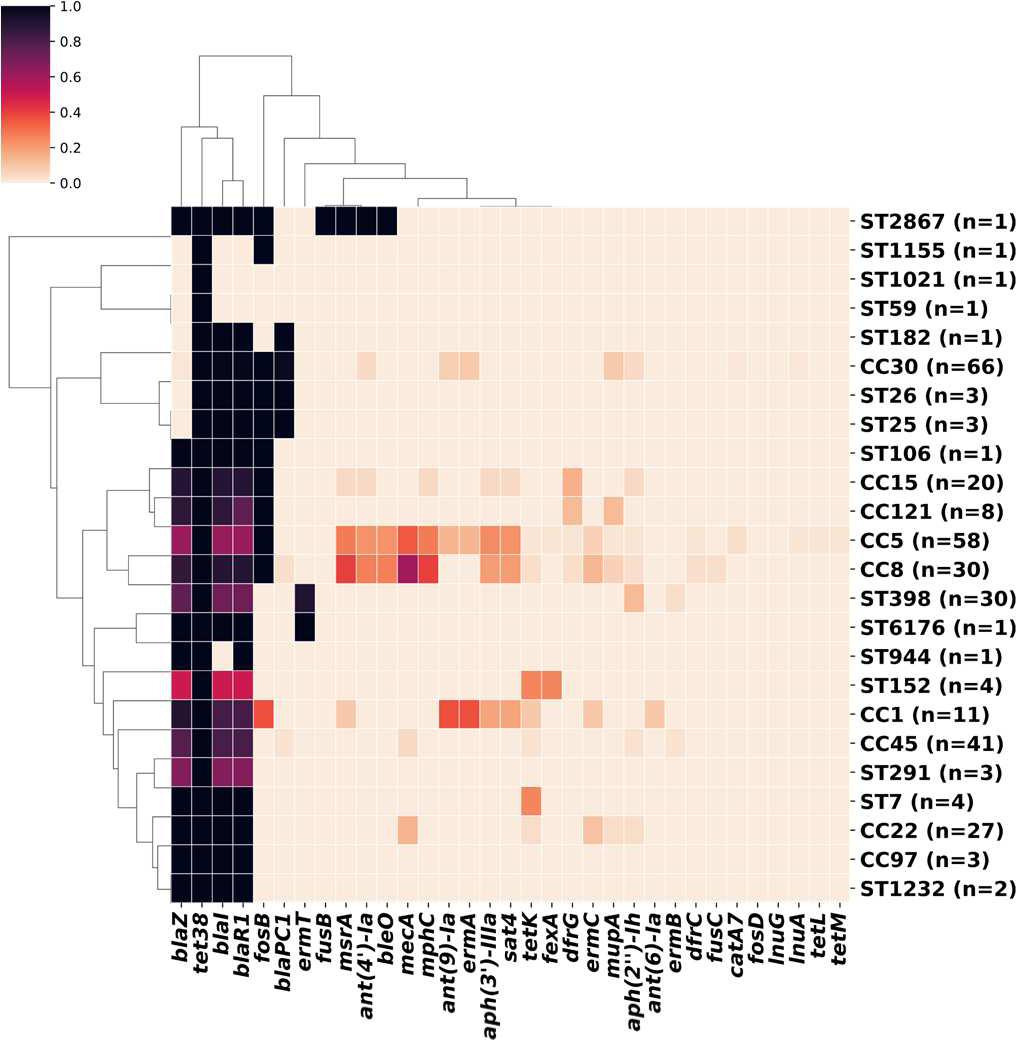
Clustered heatmap representing the frequency of abricate-predicted ARGs for all *S. aureus* lineages. Cells are colored from pale orange (0%) to black (100%). Samples were clustered and visualized using the Seaborn clustermap function. Isolates with unknown lineage were not included.

Homologs of genes related to tetracycline, β-lactams and fosfomycin resistance were the most prevalent. Homologs of *tet38* were identified in all isolates, of complete *blaI-blaR-blaZ* clusters in 279 strains (82.3%) and of *fosB* in 206 (60.2%). We found that the *blaZ_PC1_* variant was mainly associated with strains belonging to the CC30 lineage (*p* < 0.05) (Figure S1, Table S2). All isolates belonging to CC5, CC8, CC15, CC30 and CC121 lineages encoded a *fosB* gene homolog. In contrast, CC22, CC45 and ST398 clones did not present any fosfomycin resistance gene homologs (Table S2).

In addition, we identified homologs to three different groups of genes conferring macrolide resistance: several *erm* variants encoding 23S rRNA methylases (19.5%), the ABC-F type ribosomal protection encoding gene *msrA* (10.0%) and the phosphotransferase gene *mphC* (9.4%). We observed a strong correlation between homologs of *mphC* and *msrA* (MCC = 0.97), which were significatively associated with CC5 and CC8 lineages (Figure S1, Table S2). In addition, almost all strains carrying *ermT* belonged to the ST398 sequence type (*p* < 0.05) (Figure S1, Table S2). Homologs of *ermA* were associated with CC1 and CC5 while *ermC* homologs were linked to CC8 clones (*p* < 0.05) (Figure S1, Table S2). We also detected a biased distribution of the aminoglycoside nucleotidyltransferase genes *ant*(4’)-Ia (9.1%) and *ant*(9)-Ia (5.9%) as well as of the aminoglycoside phosphotransferase genes *aph*(3’)-IIIa (7.4%) and *aph*(2’’)-Ih (4.4%) (Figure S1). Specifically, *ant*(4’)-Ia gene homologs were significatively associated with CC5 and CC8 strains, *ant(9)-*Ia with CC1 and CC5, *aph(3’)-*IIIa with CC5 and CC8, and *aph(2’’)-*Ih with ST398 (Table S2). Homologs of the bleomycin resistance gene *bleO* (7.4%) showed a positive correlation with *ant*(4’)-Ia (MCC = 0.89) and were also linked to CC5 and CC8 clones (*p* < 0.05) (Figure S1, Table S2). Finally, homologs of the *mupA* gene (3.8%), conferring mupirocin resistance, were significatively associated with CC30 isolates (Figure S1, Table S2).

### Virulence factors

The presence of VF contributing to *S. aureus* pathogenicity and invasiveness was predicted (Figure 3, Data S1). All isolates from our collection carried homologs of the *cap* gene cluster responsible for the synthesis of capsular polysaccharide. Capsular polysaccharide serotype 8 was significatively more prevalent in CC15, CC30, CC45 and CC121 lineages (Figure S1, Table S3) while serotype 5 were mainly found at CC5, CC8, CC22 and ST398 clones. Homologs of the Immune Evasion Cluster (IEC) (*sak*, *chp* and *scn*) and *sbi*, also involved in immune system evasion, were present in 73.1 - 99.4% of the isolates. The *sak* gene was not found in ST398 isolates. Similarly, 99.7% of the isolates carried the *isdA-G* gene homologs, which confer resistance to killing by human lactoferrin. The intracellular adhesion (*ica*) locus, involved in cell-cell adherence and biofilm formation, was detected in all strains. Homologs of genes coding for several Microbial Surface Components Recognizing Adhesive Matrix Molecules (MSCRAMMs) were also detected, although to different extents: *ebp* (95.3%), *eap*/*map* (93.5%), *sdrC* (92.0%), *sdrE* (87.6%), *clfA* (86.1%), *fnbA* (74.6%), *clfB* (68.4%), *fnbB* (62.2%), *sdrD* (53.7%), *vWbp* (47.8%) and *cna* (15.9%). Specifically, we found that the secreted von Willebrand factor-binding protein (*vWbp*) was more prevalent in CC1, CC8, CC30, CC45 and CC121 lineages while the *cna* gene was associated with CC30 clones (*p* < 0.05) (Figure S1, Table S2).

**Figure 3.**
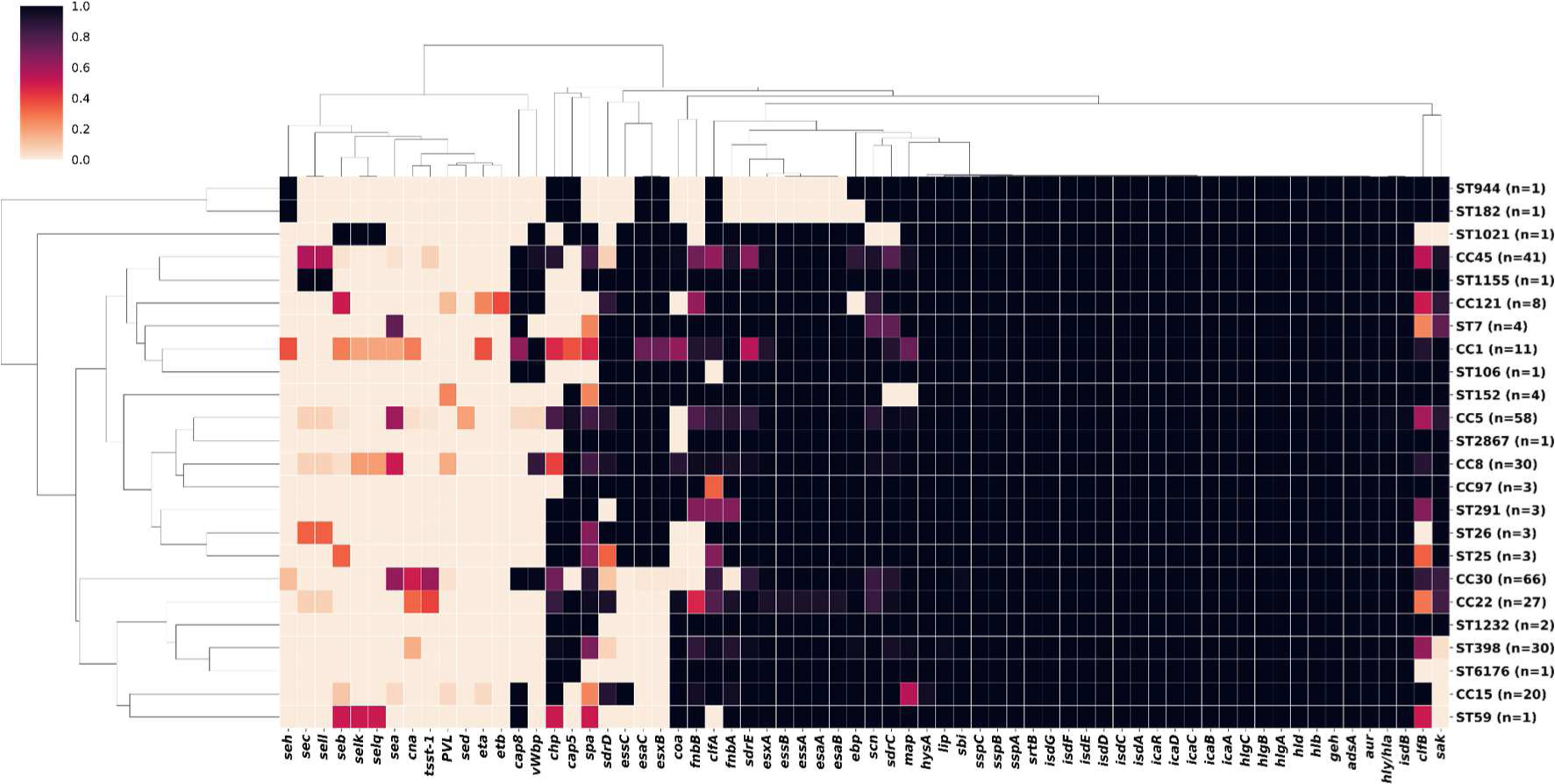
Clustered heatmap representing the frequency of abricate-predicted VFs for all *S. aureus* lineages. Cells are colored from pale orange (0%) to black (100%). Samples were clustered and visualized using the Seaborn clustermap function. Isolates with unknown lineage were not included.

Almost all isolates carried homologs of the genes encoding the α-, β- and δ-hemolysins (*hla*, *hlb*, *hlgABC* and *hld*). Genes responsible for the formation of type VII secretion systems (T7SS), *esa, ess* and *esx*, were also widespread in our collection. Enterotoxin-encoding genes (*sea, seb, sec, sed, seh, selk, sell* and *selq*) showed markedly different frequencies ranging from 31.0% (*sea*) to 2.9% (*selk* and *selq*) and significatively biased distributions among different lineages (Figure S1). Specifically, CC1 strains were associated with *seb*, *seh*, *selk* and *selq*; CC5 with *sea* and *sed*; CC8 with *sea*, *selk* and *selq*; CC30 with *sea* and *seh*; CC45 with *sea*, *sec* and *sell*; and CC121 with *seb* (*p* < 0.05) (Table S2). Of note, *sec* and *sell* genes were totally correlated (MCC = 1.0). We also detected the presence of the toxic shock syndrome toxin-1 (*tsst*-1) gene homologs in 56 isolates (16.5%), which were linked to CC22 and CC30 (*p* < 0.05) (Figure S1, Table S2). In addition, 11 isolates were predicted to encode the PVL genes (*luk*F and *lukS*) with a linkage to CC8 strains (Figure S1, Table S2). Finally, we also found homologs of the toxin-encoding genes *eta* (2.1%) and *etb* (0.9%) in our strains.

### Plasmids

We identified 51 different plasmids distributed in 256 isolates (75.5%) (Figure 4, Data S1). Among them, the vast majority (52.8%) presented a single plasmid, but we also found strains with two (18.3%), three (3.2%) or four (1.2%) plasmids.

**Figure 4.**
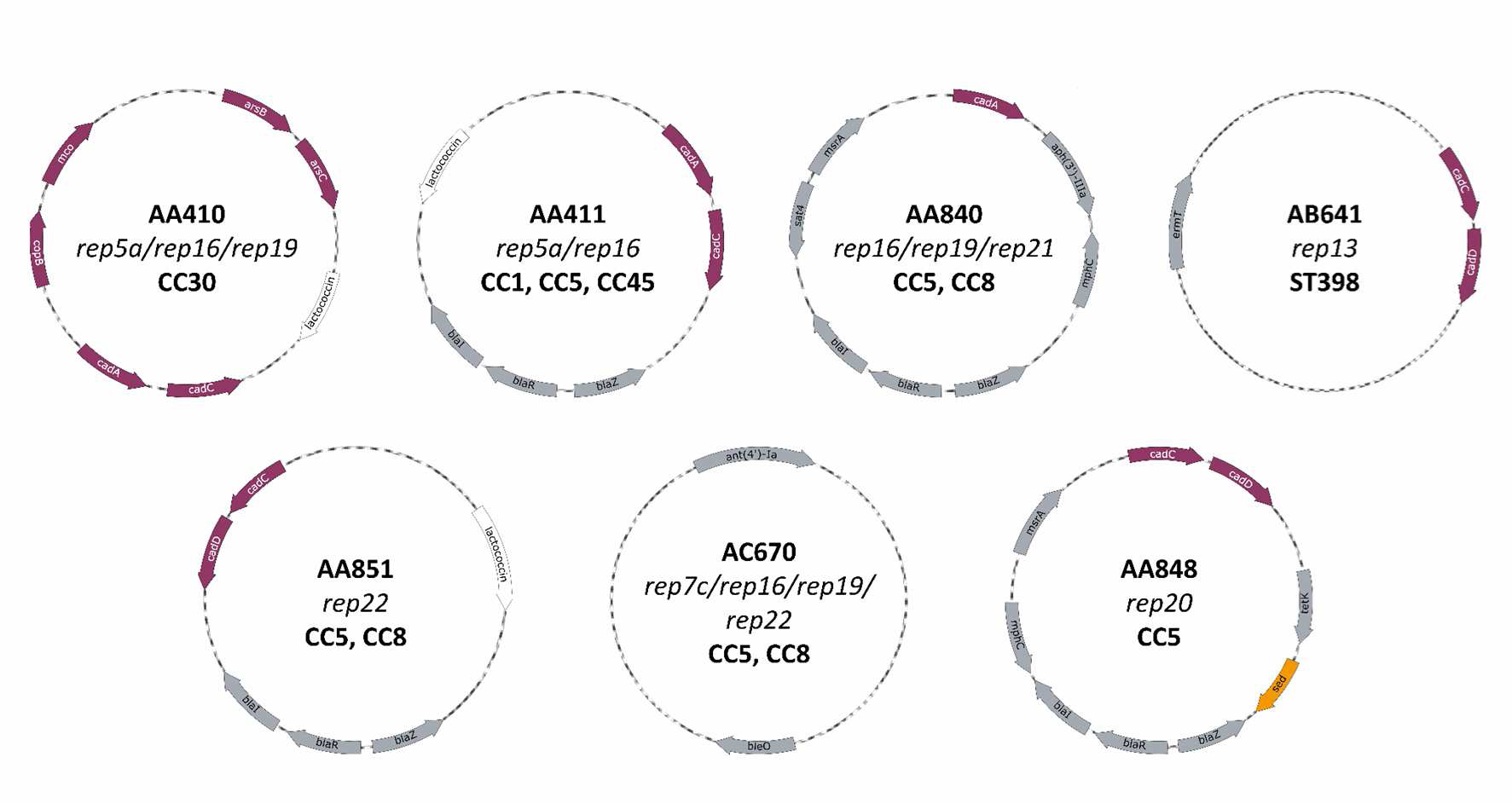
Schematic representation of the most abundant plasmids in our collection. Gene coloring depicts ARGs (gray), VF (orange), heavy metal resistance genes (purple) and lactococcin synthesis genes (white). Rep types and lineage associations are also included. Only plasmids harbored in more than 15 isolates were included.

The core proteome of the most prevalent plasmid, AA410 (n=72; *rep5a/rep16/rep19*), showed heavy metal resistance (*arsB, arsC, cadA, cadC, copB* and *mco*) and lactococcin synthesis genes (Table S3). This plasmid was linked to CC30 strains (*p* < 0.05) (Figure S1, Table S2). Plasmid AA411 (n=56; *rep5a/rep16*), harbored the *blaI-blaR-blaZ* cluster for penicillin resistance, a cadmium resistance cluster (*cadC, cadD*) and lactococcin-synthesis genes (Table S3). In this case, AA411 plasmids were significatively associated with CC1, CC15 and CC45 (Figure S1, Table S2). Plasmid AA840 (n=33; *rep16/rep19/rep21*), encoding *cadA* gene for cadmium resistance and several ARGs (*blaI-blaR-blaZ, aph(3’)-IIIa, mphC, msrA* and *sat4*) (Table S3), was more prevalent among CC5 and CC8 isolates (*p* < 0.05) (Figure S1, Table S2) and showed a positive correlation with MRSA strains (MCC = 0.73). Also significatively associated with these lineages (Figure S1, Table S2) was the AA851 plasmid (n=22, *rep22*) which harbored cadmium resistance genes (*cadC*, *cadD*), the *blaI-blaR-blaZ* cluster and genes for lactococcin synthesis (Table S3). On the other hand, the plasmid AB641 (n=25; *rep13*), which was almost exclusively detected in isolates from the ST398 lineage (*p* < 0.05) (Figure S1, Table S2), carried two cadmium resistance genes (*cadC*, *cadD*) and the macrolide-resistance gene *ermT* (Table S3). Finally, plasmid AC670 (n=19; *rep7c/rep16/rep19/rep22*) encoded genes conferring antimicrobial resistance (*ant(4’)-Ia, bleO*); and plasmid AA848 (n=15; *rep20*) carried several ARGs (*blaI-blaR-blaZ, mphC, msrA* and *tetK*), two cadmium resistance loci (*cadC*, *cadD*) and the staphylococcal enterotoxin D-encoding (*sed*) gene (Table S3). AC670 plasmids were associated with CC5 and CC8 clones while AA848 plasmids were linked to CC5 clones (*p* < 0.05) (Figure S1, Table S2).

We also predicted other less-abundant plasmids, carrying other genes conferring resistance to aminoglycosides (*ant(6)-Ia*, *aph(2’’)-Ih*), macrolides (*ermA*, *ermB, ermC, msrA, mphA*), tetracycline (*tetK*), chloramphenicol (*catA7*), lincosamides (*lnuA*) and trimethoprim (*dfrG*). The exfoliative toxin B encoding gene (*etb*) was also annotated in three plasmids (Table S3).

Overall, our results showed that *ant(6)-Ia*, *aph(3’)-IIIa, catA7, ermC, lnuA, mphC, msrA* and *sat4* genes were exclusively plasmid-borne in our collection (Figure S2). Likewise, *mupA* (92.3%), *ant(4’)-Ia* (90.3%), *bleO* (88.0%), *ermT* (80.6%) and *tetK* (77.8%) genes were also much more prevalent in plasmids than in chromosomes. Similarly, around 50% of the *blaI-blaR-blaZ* cluster was detected in plasmids. In contrast, several abundant genes such as *ant(9)-Ia, ermA, fosB, mecA* or *tet38* were always chromosomally encoded (Figure S2). Finally, the virulence genes *sed* and *etb* were found to be exclusively plasmid-borne.

### Prophages

We predicted 369 different prophage clusters in 336 isolates (97.6%) (Data S1). The majority presented one (28.6%) or two (28.9%) prophages, but we also found strains with three (21.5%), four (12.1%), five (7.4%), six (0.9%) or even seven (0.6%) prophage sequences.

Prophage clusters from the integrase group 3 (Sa3) were predominant in our collection (n=43). Some of them (n=15) carried homologs of complete IEC (*scn*, *chp*, *sak*). Enterotoxin A encoding gene (*sea*) was predicted in six of them and *tsst-1* was found in one cluster (Table S4). Of note, different Sa3-prophage clusters (1, 15, 29 and 55) showed a biased distribution in our collection (Figure S1). Specifically, cluster 1 was associated with CC30, cluster 15 with ST398, cluster 29 with CC5 and, finally, cluster 55 with CC30 and ST398 strains (*p* < 0.05) (Table S2).

We further identified some prophage clusters (n=26) of the integrase group 2 (Sa2), two of them encoding *sea* and *tsst-1* genes. We also predicted 24 prophage clusters from integrase group 1 (Sa1); 22 from integrase group 5 (Sa5), with two harboring *sec* and *sell* genes; and 17 from integrase group 6 (Sa6). Prophages from integrase group 7 (Sa7) and 9 (Sa9) were also identified but to a lesser extent. Finally, 221 prophage clusters did not present any integrase gene. Among those, 14 clusters carried a complete IEC and 10 harbored the *sea* gene (Table S4).

No ARG were detected in the predicted prophage genomes.

### Pandemic-associated features

We identified 139 and 200 consecutive adult patients diagnosed with bloodstream infections by *S. aureus* before and after March 14th 2020, respectively (Data S1). In the pre-pandemic years, we detected 27.8 episodes of *S. aureus* bacteremia per year in our hospital rising to 66.7 during the pandemic. In such years, we observed an increase of nosocomial infections, from 69 episodes in the pre-pandemic years to 119 during the COVID-19 pandemic. Specifically, cases of *S. aureus* peripheral catheter-related bacteremia were significantly more prevalent and we also found 2.5 times more episodes of bacteremia originating from surgical sites (Table S5). We also identified a significant increase of MRSA infections (*p* < 0.05) and we diagnosed 2.8 times more endocarditis cases (Table S5).

Statistical analysis on pre- and pandemic strains revealed significant genetic differences between these two *S. aureus* populations. We observed a strong increase of strains carrying genes encoding the clumping factors A (*clfA*, *p* = 2.1e^-03^) and B (*clfB, p* = 2.3e^-05^) (Table S5). Clumping factors A and B are MSCRAMMs that bind fibrinogen, which is found at high levels upon SARS-CoV-2 infection (O’Brien et al. 2002; Kangro et al. 2022). In contrast, we found a significant reduction of strains encoding the *sdrE* gene, which is also classified as MSCRAMM (Table S5). On the other hand, we noticed a rise in several antibiotic resistance genes during those years. Specifically, the macrolide-resistance *msrA* and *mphC* genes as well as the methicillin-resistance *mecA gene* showed a significant increase during the pandemic years (Table S5). Of note, macrolides were extensively used as accessory therapy for COVID-19 in patients (Sterenczak et al. 2020; Grau et al. 2021).

## DISCUSSION

### Global insights into *S. aureus* clinics, antibiotic resistance and virulence

Genomics enables unraveling the genetic makeup of pathogens, providing insights into virulence factors, antibiotic resistance mechanisms and the overall pathogenic potential of bacteria. This information is crucial for the development of targeted diagnostics, treatment strategies and preventive measures. The use of genomics also allows tracking the spread of specific strains and identifying emerging threats, contributing to the surveillance and control of infections. Here, using a collection of 339 *S. aureus* isolates from patients with bacteremia, we unveiled important insights into the clinical and genomic features of these bacterial pathogens (Table 1).

**Table 1.** Summary of the significant associations between the most abundant *S. aureus* lineages with all the clinical and genetic data.

| Lineage | Clinical features | Gene types | Antibiotic Resistance Genes | Virulence Factors | Plasmids | Prophages |
| --- | --- | --- | --- | --- | --- | --- |
| CC1 |  |  | *ant(9)-Ia, ermA* | *seb, seh, selk, selq* | AA411 |  |
| CC5 | MRSA, pneumonia-originated bacteremia | SCCmec IVc, *agr* type II, *spa* t002, *spa* t067 | *ant(4')-Ia, ant(9)-Ia, aph(3')-IIIa, bleO, ermA, fosB, mecA, mphC, msrA* | *cap5, sea, sed* | AA840, AA851, AC670, AA848 | cluster_29 (Sa3) |
| CC8 | MRSA, pneumonia-originated bacteremia | SCCmec IVc, *spa* t008 | *ant(4')-Ia, aph(3')-IIIa, bleO, ermC, fosB, mecA, mphC, msrA* | *cap5, sea, selk, selq, PVL* | AA840, AA851, AC670 |  |
| CC15 |  | *agr* type II, *spa* t084 | *fosB* | *cap8* | AA411 |  |
| CC22 |  | SCCmec IV |  | *cap5, tsst-1* |  |  |
| CC30 | Bacteremia from surgical site infections | *agr* type III, *spa* t012, | *blaZPC1, fosB, mupA* | *cap8, sea, seh, tsst-1* | AA410 | cluster_1 (Sa3), cluster_55 (Sa3) |
| CC45 |  |  |  | *cap8, sea, sec, sell* | AA411 |  |
| CC121 |  |  | *fosB* | *cap8, seb* |  |  |
| ST398 |  | *spa* t1451 | *aph(2'')-Ih, ermT* | *cap5* | AB641 | cluster_15 (Sa3), cluster_55 (Sa3) |

In our hospital, we identified *S. aureus* lineages mirroring those previously reported in Spain (Pérez-Montarelo et al. 2018) and in other countries (Fowler et al. 2007; Park et al. 2017; Campbell et al. 2022) (Figure 1), suggesting a shared genetic heritage and potentially common evolutionary pathways. Pérez-Montarelo and colleagues analyzed strains isolated across various Spanish hospitals spanning 2002 to 2017 (Pérez-Montarelo et al. 2018). Our study, conducted between 2014 and 2022, complements this report and broadens its scope into the pandemic period, offering compelling observations on the evolutionary trends of *S. aureus* clones. Specifically, we observed an increase of ST398 lineages, heightening from 2.4% (Pérez-Montarelo et al. 2018) to a mean prevalence of 8.8% in our isolates. Importantly, the prevalence of this lineage in our collection ranged from 0% (2014 and 2015) to 15% (2022). These results highlight the spread of ST398 lineage in hospital settings, confirming the need to focus on the mechanisms involved in the epidemiological success of ST398 in humans, as previously suggested (Uhlemann et al. 2017; Mama et al. 2021; van der Mee-Marquet et al. 2022) (Figure 1).

Despite possible associations between genotypes and certain complications, the role of *S. aureus* genetic factors in the evolution of bacteremia is still poorly understood. CC1, CC5, CC8, CC22 and CC30 lineages have been associated with increased risk of complications (Fowler et al. 2007; Miller et al. 2012). In contrast, more recent studies did not find any apparent association between clonality and poorer outcomes (Chong et al. 2013; Gasch et al. 2014; Fernández-Hidalgo et al. 2018). Our study is in line with the latter since we did not find significant associations between the different isolates from our collection and clinical outcome. However, there is a tendency for the CC1 and CC8 strains to cause fatal infections. On the other hand, we identified associations between the different lineages and the focus of infection. CC5 and CC8 were correlated with pneumonia-originated bacteriemia, as deduced from previous reports (Antonelli et al. 2019), and CC30 with surgical site infections (Table 1, Table S2).

MRSA poses a significant public health concern due to its ability to cause infections that are challenging to treat, showing higher mortality than MSSA infections (Munckhof et al. 2008). In Spain, the prevalence of methicillin resistance among *S. aureus* isolates is around 20-25% (Pérez-Montarelo et al. 2018; Vázquez-Sánchez et al. 2022), while the incidence of MRSA infections in our hospital showed to be 13.9% (Figure 2). Of note, we did not find associations with mortality and MRSA strains. In accordance with some reports, most of our MRSA isolates belonged to CC5 or CC8 strains (Dupper et al. 2019). These lineages were also the ones accumulating more antibiotic resistance genes (Figure 2, Table 1), as previously stated (Smith et al. 2021). Genes predicted to confer resistance to tetracycline and β-lactams were the most prevalent in our isolates (Figure 2). However, the results for tetracycline did not align well with the patterns observed *in vitro* by broth microdilution tests. The *tet38* gene, which was present in almost all strains, is known to confer tetracycline resistance when overexpressed (Chen and Hooper 2018), but its mere presence does not necessarily indicate resistance to this class of antibiotics.

*S. aureus* is known for its pathogenic potential, primarily attributed to a diverse set of virulence factors. As previously elucidated, the *agr* groups and capsular polysaccharide serotypes were associated with *S. aureus* clonality (Figure 3, Table 1) (Monecke et al. 2011). Several genes involved in immune system evasion (*adsA*, *aur*, *sbi, ssl, sspA, sspB*), lactoferrin resistance (*isdA-G*), adhesion (*ica*), T7SS formation (*esa, ess, esx*), some MSCRAMMs (*ebp*, *eap/map*, *sdrC*) and hemolysins were found in virtually all strains (Figure 3), similar to previous reports (Rasmussen et al. 2013). IEC is an immune response cluster harboring *chp*, *scn* and *sak* genes which is typically encoded by β-hemolysin converting bacteriophages (Sieber et al. 2020). This is partially paralleled in our isolates, in which we predicted 61.7% of complete IEC clusters in prophage regions (Table S4). In contrast to many other bacterial pathogens, *S. aureus* produces a wide array of toxins (Cheung et al. 2021). In our hospital, besides hemolysins, the most abundant predicted toxins were the exotoxins SEA (31.0%) and TSST-1 (16.5%), the latter being the major cause of toxic shock syndrome (Figure 3). SEA is the most common staphylococcal enterotoxin (Pinchuk et al. 2010) and is also reported to be encoded by a prophage (Betley and Mekalanos 1985). In our isolates, only 53.8% of *sea* genes were predicted in prophage regions (Table S4), hereby indicating that we could be underestimating prophage regions. Finally, we found 11 PVL-positive strains (3.2%) (Figure 3). Panton–Valentine leukocidin is a pore-forming leukotoxin produced by 2-3% of *S. aureus* isolates (Wójcik-Bojek et al. 2022) which has been traditionally used as an indicator of community-acquired MRSA strains (Fey et al. 2003). However, some recent studies found that PVL may not be a reliable marker because it is being reported in nosocomial MRSA infections (Nakaminami et al. 2020; Shohayeb et al. 2023). Paralleling these reports, we found 4 non-community MRSA harboring PVL suggesting a possible spread of these strains in hospitals.

### Changes in molecular characteristics of pandemic *S. aureus* isolates

Understanding the interplay between viral and bacterial infections, including the prevalence and impact of *S. aureus* in COVID-19 cases, is important for effective patient care and public health measures. Several studies underscored a higher incidence of bacteremia by *S. aureus* at the COVID-19 pandemic (Falces-Romero et al. 2023). In keeping with this, the episodes of *S. aureus* bacteremia per year doubled in our hospital during the pandemic. Concomitantly, we found a rise in nosocomial infections, particularly those originated from catheters and surgical sites. The increase of catheter-related bacteremia during the COVID-19 epidemic has been previously documented by our group (Gasch et al. 2022).

Unlike other countries, we did not detect emerging *S. aureus* clones during the pandemic (Gu et al. 2023). However, we noticed a significant increase in strains carrying the *clfA* and *clfB* genes, a reduction of strains encoding *sdrE*, along with a rise in macrolide and methicillin antibiotic resistance genes (Table S5).

The *clfA* and *clfB* genes are MSCRAMMs encoding the fibrinogen-binding Clumping Factors A and B, respectively. Fibrinogen is one of the most prevalent coagulation proteins in blood (Kangro et al. 2022). ClfA is the major virulence factor responsible for *S. aureus* clumping in blood plasma (O’Brien et al. 2002) and ClfB interacts with cytokeratin 10 and loricrin facilitating *S. aureus* skin infection (Lacey et al. 2019). Both ClfA and ClfB have been reported to be involved in bacterial and platelet aggregation (O’Brien et al. 2002). Interestingly, several reports documented an increase of fibrinogen production by SARS-CoV-2 virus (Bouck et al. 2021; Kangro et al. 2022). Therefore, our results suggest that the presence of *clfA* and *clfB* genes provided an advantage to *S. aureus* in the context of SARS-CoV-2 co-infection by enhancing fibrinogen binding. In consequence, *S. aureus* bloodstream isolates carrying the *clfA* and *clfB* genes were positively selected between 2020 and 2022. Conversely, we found a reduced prevalence of the *sdrE* gene, which codes for a MSCRAMM that prevents *S. aureus* phagocytosis by sequestering the human complement factor H (CFH) on the surface of bacterial cells (Zhang et al. 2017; Wójcik-Bojek et al. 2022). This is remarkable because it has been shown that the SARS-CoV-2 spike protein blocks CFH leading to a complement dysregulation on the human cell surface (Yu et al. 2022). Therefore, the functional redundancy of *S. aureus* SdrE and the SARS-CoV-2 spike protein could explain the negative selection of *sdrE*. In the same line, it is important to note that *S. aureus* CflA has also been found to have an anti-phagocytic effect (Higgins et al. 2006; Hair et al. 2010).

Regarding *S. aureus* antibiotic resistance, some articles have emphasized the role of macrolides as adjunctive therapy for COVID-19 in certain patients (Sterenczak et al. 2020) and the use of such antibiotics could allow selecting resistant strains. Macrolide resistance in staphylococci is mainly associated with mobile ARGs which often correlates with methicillin resistance (Serra et al. 2023), as also observed in our collection (Figure 2). These findings emphasize the evolving nature and impact of *S. aureus* infections in the COVID-19 pandemic era.

## FUNDING

This work was supported by the grant PI19/01911 from Instituto de Salud Carlos III (ISCIII). MSO and PBC were the recipients of a Margarita Salas fellowship from the Ministerio de Universidades and a PFIS fellowship from the Instituto de Salud Carlos III, respectively.

## ACKNOWLEDGMENTS

We are grateful to the staff of the Genomics Unit (CRG) for Illumina sequencing. We also want to acknowledge the CERCA Programme / Generalitat de Catalunya.

## Author Contributions

Conceptualization, MS-O and OQP; Methodology, MS-O, IE, OG and OQP; Software, MS-O and IE; Investigation, MS-O, MP, PB, IG-S, MA-G, ME, IE, OG and OQP; Data Curation, MS-O and MP; Writing—Original Draft Preparation, MS-O and OQP; Writing—Review and Editing, MS-O, MP, PB, IG-S, MA-G, ME, IE, OG and OQP; Funding Acquisition, OQP. All authors have read and agreed to the published version of the manuscript.

## CONFLICT OF INTEREST STATEMENT

The authors declare that the research was conducted in the absence of any financial or commercial relationships that could be a potential conflict of interest.

## DATA AVAILABILITY

Sequence data generated in this study have been deposited in the NCBI database with the following access number: PRJNA1055690. Source data are provided with this paper as Supplementary Material.

## SUPPLEMENTARY MATERIAL

**Data S1.** JSON-formatted file including the clinical metadata and predicted features for all *S. aureus* strains included in this study.

**Figure S1.** Pie plots depicting lineage frequencies for all clinical and genetic features. The MWU *p*-values obtained when comparing to global lineage frequency are included.

**Figure S2.** Heatmap representing the frequency of chromosome- or plasmid-encoded genes for all ARGs in *S. aureus* lineages. Cells are colored from blue (all instances are plasmid-borne) to red (all instances are in the chromosome), with white indicating absence of ARG.

**Table S1.** Results for statistical analysis of clinical data. In each case, the computed *p*-value and selected statistical method are included.

**Table S2.** Results of Fisher’s exact test (FT) for analyzing associations between specific *S. aureus* lineages and clinical or genetic data. FT was only computed for features with biased distributions in MWU analysis (Figure S1).

**Table S3.** List of predicted plasmids including number of instances, the roary-core proteome, and the abricate-predicted antibiotic resistance genes and virulence factors.

**Table S4.** List of predicted prophage clusters including number of instances, integrase type, presence of IEC and the abricate-predicted virulence factors. All phage instances of the same cluster not necessarily harbored all the features.

**Table S5.** Results of Fisher’s exact test for comparing the clinical and genetic features of pre- and pandemic bacteremia episodes.

## Notes

### Competing Interest Statement

The authors have declared no competing interest.

